# Optical-flow analysis toolbox for characterization of spatiotemporal dynamics in mesoscale optical imaging of brain activity

**DOI:** 10.1101/087676

**Authors:** Navvab Afrashteh, Samsoon Inayat, Mostafa Mohsenvand, Majid H. Mohajerani

## Abstract

Wide-field optical imaging techniques constitute powerful tools to sample and study mesoscale neuronal activity. The sampled data constitutes a sequence of image frames in which one can perceive the flow of brain activity starting and terminating at source and sink locations respectively. The most common data analyses include qualitative assessment to identify sources and sinks of activity as well as their trajectories. The quantitative analyses is mostly based on computing the temporal variation of the intensity of pixels while a few studies have also reported estimates of wave motion using optical-flow techniques from computer vision. A comprehensive toolbox for the quantitative analyses of mesoscale brain activity data however is still missing. We present a graphical-user-interface based Matlab^®^ toolbox for investigating the spatiotemporal dynamics of mesoscale brain activity using optical-flow analyses. The toolbox includes the implementation of three optical-flow methods namely Horn-Schunck, Combined Local-Global, and Temporospatial algorithms for estimating velocity vector fields of perceived flow in mesoscale brain activity. From the velocity vector fields we determine the locations of sources and sinks as well as the trajectories and temporal velocities of activity flow. Using our toolbox, we compare the efficacy of the three optical-flow methods for determining spatiotemporal dynamics by using simulated data. We also demonstrate the application of optical-flow methods onto sensory-evoked calcium and voltage imaging data. Our results indicate that the combined local-global method we employ, yields results that correlate with the manual assessment. The automated approach permits rapid and effective quantification of mesoscale brain dynamics and may facilitate the study of brain function in response to new experiences or pathology.

**Conflicts of Interest:** none

**Author contribution statement:** MHM, MM, NV, and SI designed the study. NA and SI wrote Matlab^®^ code for the toolbox and designed the simulated data. MHM, and NA performed the experiments. NA and SI analyzed the data. SI, NA, and MHM wrote the manuscript.

## 1. Introduction

In order to investigate and build physiological models of brain functions at various levels of abstraction *viz* molecular, cellular, system, and behavioral levels, evoked or spontaneous activity of the brain is observed at different spatial scales namely nano, micro, meso, macro, and eco. At the mesoscale corresponding to spatial dimensions ranging from hundreds of micrometers to a few centimeters, activity of thousands of neurons can be recorded simultaneously to study how networks of neurons communicate with each other. Various methods have been developed to monitor mesoscale brain activity such as intrinsic or extrinsic optical imaging, and multielectrode electrophysiology (1–5). In optical imaging, brain is excited with light and imaged to detect neuronal activity at the population level as changes in the intensity of captured light. The neuronal activity is transduced into an optical signal by either intrinsic or extrinsic sensors (4,6–8). In imaging with intrinsic sensors such as flavoprotein fluorescence imaging, endogenous autofluorescence of molecules representing biological activity is detected. In imaging with extrinsic sensors however, fluorescent molecules are introduced into the brain and depending upon their chemical structure, the fluorescence changes are modulated by either neuronal membrane voltage (voltage imaging), or concentration of an intracellular ion (e.g. calcium imaging) or molecule (9–13). Thus the data obtained with optical imaging is a set of images where the intensity of each pixel is representative of the summed activity of nearby neurons. Similarly, in multielectrode electrophysiology, electrodes are spread over surface of the brain and enable spatially discrete sampling of neuronal activity. The electrophysiology data can also be visualized as a set of 2D images where the intensity of each pixel would represent the voltage measured by an electrode (14). Mesoscale sampling of brain activity hence captures the spatiotemporal dynamics of networks of neurons resulting in data with one temporal and two spatial dimensions (Fig. 1).

**Figure 1.**
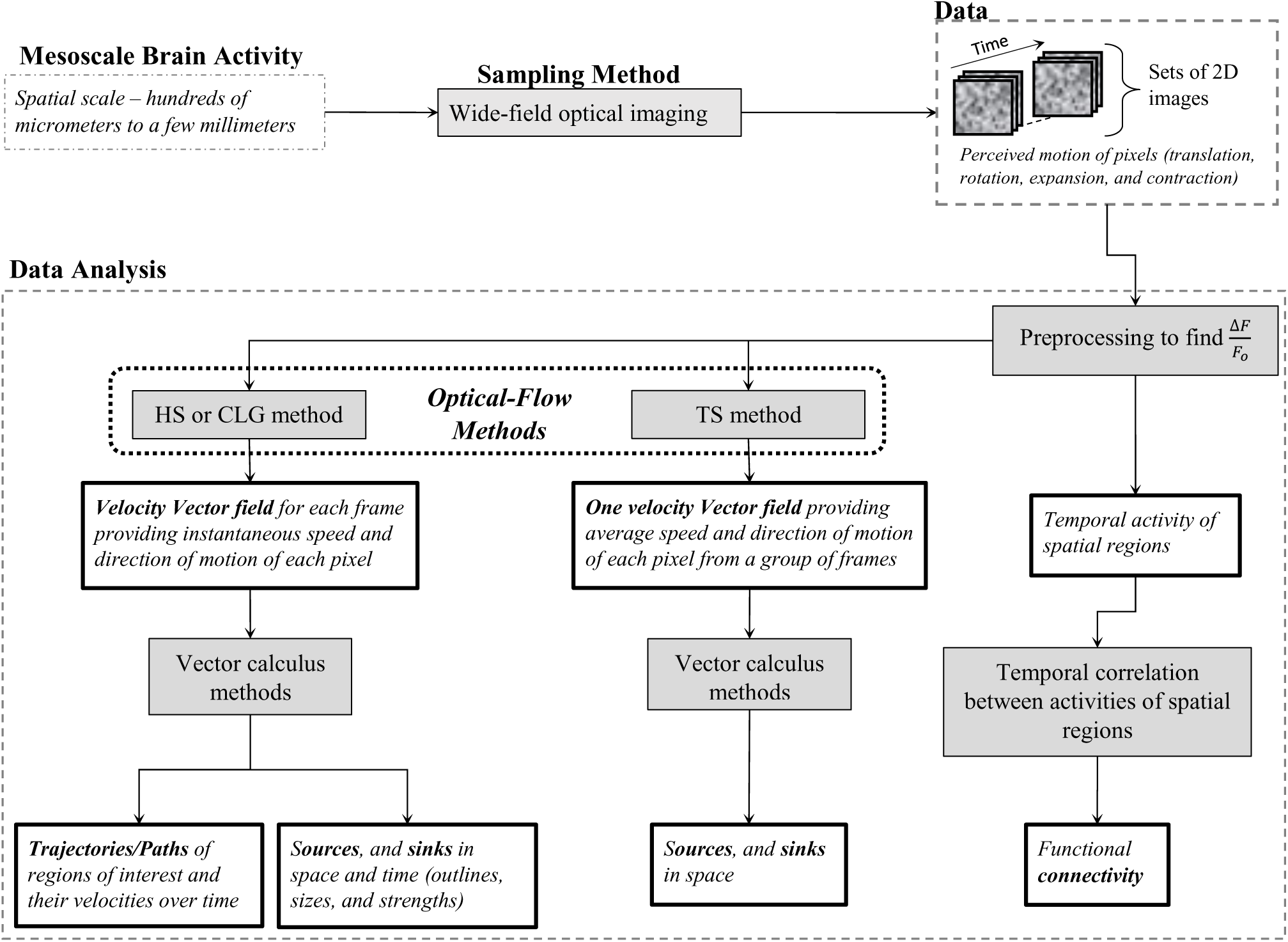
Flow diagram summarizing mesoscale brain activity data acquisition and analysis. The data consists of perceived motion of pixels i.e. traveling waves originating at sources and vanishing at sink locations. The most common analysis includes observing temporal activity of pixels or regions of interest and determining functional connectivity between spatial regions using temporal correlations. The less common analysis is estimating wave motion with “optical-flow methods” (borrowed from computer vision) in which the spatiotemporal dynamics in brain activity are characterized for determining source/sink locations and properties of traveling waves i.e. trajectories and temporal speeds of pixels. We compare the performance of three optical flow methods; Horn-Schunk (HS), Combined local-global (CLG), and Temporospatial (TS) for analyzing optical imaging data. We present a graphical user interface based Matlab toolbox for mesoscale brain activity data analysis. Shaded boxes show methods included in the toolbox and bold outlined boxes show output entities of interest.

There are a variety of features observed in optical imaging of brain activity. For example, if a group of neurons become active spontaneously or in response to a stimulus, a source like activity is observed where pixel intensities gradually increase. The observed shape of the source-like activity in 2D might be a point, line, circle, ellipse, or a combination of them depending upon the anatomical organization of active neurons. The source activity, later in time, might disappear or sink in the same spatial region or travel like a wave to another region following a path/trajectory and then disappear into a sink where pixel intensities gradually decrease. The sink might also have shapes in 2D similar to the source activity. For propagating activity, the paths might be a mix of translational, rotational, expansion or compression trajectories. If multiple brain areas are simultaneously active, multiple sources, travelling waves, and sinks would be observable. The observation of brain activity at the mesoscale hence mimics the motion or flow of waves.

The most common analysis used for the extraction of physiological features from optical imaging of brain activity is the quantitative assessment of temporal variation of intensity by plotting the average intensity of pixels versus time for defined regions of interest. More recently, the temporal correlations of optical signals were used to determine the functional connectivity between different regions (Fig. 1) (15–21). To identify sources, sinks, and activity trajectories manual methods were often used (16,22). However, few studies have reported estimates of neuronal activity spread using techniques first developed in the field of computer vision referred to as “optical-flow methods”. In these methods, velocity vector flow fields are calculated to determine speeds and directions of motion (23,24). From the velocity vector fields, locations of sources and sinks are estimated using vector calculus methods. Inouye et al. 1994 (25–27) reported the use of Horn-Schunk (HS) method (28) to determine the flow of brain oscillations over the human scalp in flattened (3D surface to 2D) electroencephalography data. Takagaki et al. 2011 (29) developed and reported a new method (Temporospatial – TS) in which temporal correlation of a given pixel with neighboring pixels was used to estimate local motion in voltage sensitive dye (VSD) imaging data. Recently, Mohajerani et al. 2013 (16) used the combined local-global (CLG) algorithm (30,31) to determine velocity vector fields of flow in wide-field VSD imaging data. They also manually determined the location of sources and sinks in space and time. Following Mohajerani et al., Townsend et al. 2015 (3) also used the CLG method to estimate wave patterns in cortical activity recorded with multielectrode arrays.

Only HS, CLG, and TS optical flow methods have been used for the analysis of brain activity but there are numerous other optical flow algorithms that have been developed (32,23,24,33–35). In spite of the above-mentioned attempts to estimate wave motion in brain activity, it is still pending to compare the performance and analytical efficacy of optical-flow methods for determining velocity vector fields in brain activity. There also has been a scarcity of quantitative analysis for extracting spatiotemporal dynamics in brain activity from velocity vector fields. Off the shelf tools are also missing for streamlining data analysis i.e. estimating velocity vector fields followed by automatically identifying sources and sinks, and calculating trajectories and velocities of brain activity with respect to time. In addition to identifying sources and sinks in space-time, it would also be useful to quantitatively characterize their properties such as outlines (what is the shape of a source or a sink), sizes (how big a source or sink is in space) and strengths (how much activity outflow or inflow is there in time).

In this paper, we present for the neuroscience community, a graphical user interface based ***O**ptical-**F**low **A**nalysis Toolbox in **M**atlab*^®^ (Mathworks Inc.) for investigating the spatiotemporal dynamics of ***m**esoscale brain activity* (**OFAMM**). We compare the performance and analytical efficacy of three optical-flow methods namely, Horn-Schunck (HS), combined local-global (CLG), and temporospatial (TS) for determining velocity vector fields of perceived flow in brain activity monitored using voltage and calcium imaging. Also, we compare the performance of HS and CLG methods based analyses in determining sources and sinks in space-time as well as trajectories of brain activity waves and their temporal speeds. The three optical flow methods were selected from numerous others (23), primarily, due to their previous use with brain activity. Since in real data sets of brain activity, the ground true values are unknown, simulated data was first used to investigate the accuracy of our analysis and its sensitivity to the addition of noise. Later, we tested and validated the application of our analysis on real data acquired with wide-field voltage and calcium imaging from mice brains. Similar to previous finding, higher instantaneous and temporal speeds were estimated with voltage imaging as compared to calcium imaging. The results are consistent with voltage signals reporting predominantly subthreshold activity and calcium signals reporting suprathreshold spiking activity (36).

## Materials and Methods

### 2.1 The General framework

The ultimate goal of analyzing the sampled brain activity with optical-flow methods is to characterize and study the perceived motion of activity (in space-time). To do so, the set of two-dimensional (2D) images collected over time are first preprocessed to filter noise and determine the percentage change in fluorescence from a baseline (ΔF/F_o_) for each pixel. Next using optical-flow methods “velocity vector fields” are determined by estimating the displacement of pixels over time. Thus, a velocity vector is estimated for each pixel whose magnitude and direction represents its speed and direction of motion. Here, we used three different optical flow methods HS, CLG, and TS for determining the velocity vector fields (Fig. 1). With the TS method, only one vector field is estimated for a group of image frames whereas with the CLG and HS methods vector fields are obtained for all pairs of consecutive image frames. From the vector fields, the locations of sources and sinks i.e. the regions of origin and termination of activity, can be determined using vector calculus methods. Since with the TS method, only one vector field is estimated for a group of frames, the location of sources and sinks can be estimated only in space but not in time (29). However, from the vector fields estimated by the CLG and HS methods the location of sources and sinks can be estimated in both space and time and more importantly, the trajectories of pixels or regions of interest and their temporal velocities can also be calculated using vector calculus methods (Fig. 1). The details of the optical-flow and vector calculus methods that we use in this work are given below.

### 2.2 Optical-flow estimation methods

#### 2.2.1 Horn-Schunck (HS) method

The HS method (28,37) operates on two consecutive frames and estimates the motion of pixels from one frame to the other by iteratively solving an optimization problem (stated below) formulated from two constraints. The first constraint called “brightness constancy” assumes a pixel to have the same brightness level in both frames after movement i.e.

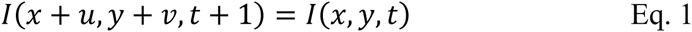

where *I(x,y,t)* is the pixel brightness in the first frame at time t and spatial location *(x,y)* and *I(x+u,y+v,t+1)* is the pixel brightness in the second frame at t+1 after *(u,v)* displacements from *(x,y)* in x and y directions respectively. The second constraint called “spatial smoothness” prevents discontinuities in the flow field. Mathematically,

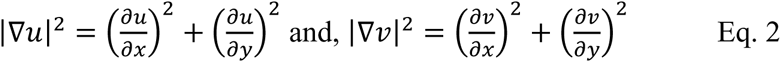

These two constraints are combined to get a minimization problem given by the following equation

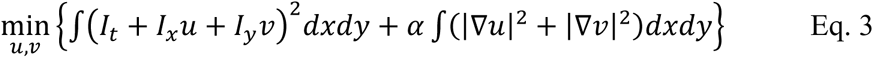

where *I_t_*, *I_x_*, and *I_y_* are derivatives of *I(x,y,t)* with respect to time, spatial direction *x*, and spatial direction *y*, respectively (17) and α is the ratio of the weights of spatial smoothness integral to that of the brightness constancy. This constant regulates the contribution of the two terms. This equation is solved to estimate the values of *u* and *v*. As the integral in the above equation is calculated on the whole image field this method is called a global method. An implementation of the HS method in Matlab^®^ was downloaded from (<u>https://www.mathworks.com/matlabcentral/fileexchange/22756-horn-schunck-optical-flow-method</u>) and used here for analysis. α, and the number of iterations can be set in the graphical-user-interface of the toolbox. We chose α = 0.1 and the number of iterations = 2000 for our subsequent analysis by trial and error. We estimated velocities of pixels with the HS method for the travelling plane and circular waves (simulated data) while tweaking parameter values to minimize deviation from actual velocities. Alternately, α and the number of iterations were also determined analytically by using simulated data with known values of trajectories and velocities for the three Gaussian waves (see Supplementary Movie 1 – last clip). Different combinations of the values of α (ranging between 0.05 and 100), and the number of iterations (ranging between 100 and 9000) were used to estimate the best combination that gave the minimum error for estimated velocities. The best combination of α = 0.35 and number of iterations = 2000 was used for analysis for generating supplementary figures S3-S7.

#### 2.2.2 Combined local-global (CLG) method

The CLG method (30,31) combines the HS method and a second method called Lucas-Kanade (LK). LK method assumes the constancy of motion in the neighborhood of a pixel i.e. the velocity and direction of motion of pixels around the pixel of interest are equal. Mathematically, LK method formulates the following minimization problem

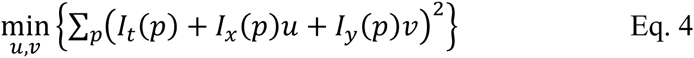

where *p* is selected from an *n* × *n* neighborhood of (*x*, *y*) pixel. In a more general formulation of the LK method the optimization argument can be convolved with a Gaussian window to filter noise in the images i.e.

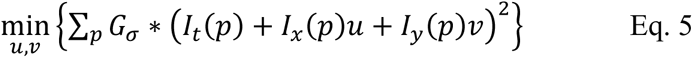

where *G*_*σ*_ is a two dimensional normalized Gaussian window with standard deviation of *σ* and ∗ denote the convolution operation. As the LK method assumes that the motion (speed and direction) is constant in the neighborhood of a pixel, it is called a local method and works better while dealing with small displacements.

The combination of HS and LK methods proposed by Bruhn et al. 2002 (30) constitutes the CLG method and is formulated as bellow:

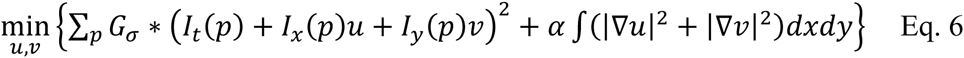

An implementation of the CLG method in Matlab (38) was downloaded from (<u>http://people.csail.mit.edu/celiu/OpticalFlow/</u>) and used here for analysis. The values of the following parameters can be set via the graphical-user-interface of the toolbox. We chose the values indicated below following (38), and trial and error in order to minimize the error in the estimation of velocities for the travelling plane and circular waves (simulated data).

*α* = 0.03, *ratio* = 0.5, *minWidth* = *Image Size* × *ratio* × 0.5, *nOuterFPIterations* = 7, *nInnerFPIterations* = 1, *nSORIterations* = 30

Alternately, the following values were used for generating supplementary figures S3-S7.

*α* = 0.04, *ratio* = 0.5, *minWidth* = *Image Size* × *ratio* × 0.5, *nOuterFPIterations* = 15, *nInnerFPIterations* = 3, *nSORIterations* = 30

These values were obtained analytically by using trial and error application of CLG method onto simulated data same as that used for the HS method (see Supplementary Movie 1 – last clip). First, a large number of iterations was used and the values of *α* and ratio were determined which gave the minimum error in the estimated velocities. Later, the number of iterations were reduced to a desired number which kept the same error to a minimum.

#### 2.2.3 Temporospatial (TS) method

The TS method proposed and demonstrated by Takagaki et al. 2011 (29) estimates motion of pixels by identifying neighboring pixels whose temporal activity is highly correlated and hence was named Temporospatial method. To determine the speed and direction of motion of a pixel, the correlation of its activity with that of neighboring pixels is calculated to determine the highest correlation coefficient and thus estimate the next location of the pixel. The speed of motion is then determined by dividing the displacement of pixel from one location to the other by the best temporal lag (delay) that provides the highest correlation coefficient. Finding temporal correlation of all pairs of pixels is a computationally intensive process. To reduce computational time, Takagaki et al. defined spatial motion templates of pixels which represented different types of motion (e.g. expansion, contraction, translation, and rotation) as bases along which the motion of pixels could be resolved. The motion of a pixel was then estimated by finding the components of its motion along the bases (see Takagaki et al. 2011 for details). Since the whole temporal activity of a pixel in a group of frames is used to determine temporal correlations with neighboring pixels, only one vector is obtained for a pixel representing the average movement of that pixel in time. Thus for a group of image frames only one vector field is obtained. The implementation of TS method we used in this work was written in Matlab^®^.

### 2.3 Vector calculus methods

#### 2.3.1 Determination of the locations of sources and sinks from velocity vector fields

The locations of sources and sinks (points of outward or inward flow respectively) from velocity vector fields were calculated using three methods. In the first method, divergence of the vector field was calculated to obtain a scalar field with positive and negative numbers indicating locations of sources and sinks respectively while their magnitude indicates the strength. Contours of constant divergence values were also calculated. In the second method, Poincare index was calculated for a pixel by finding the sum of differences of angles of neighboring pixels. Poincare index when normalized by 2π, has values of +1, 0, and −1 for a source or a sink, plane flow, and saddle points, respectively. In the third method, trace and determinant values of the Jacobian matrix were calculated for each pixel. A pixel is a source if the determinant and trace values are positive and is a sink if the determinant is positive with a negative trace value. A pixel was only considered a source (or a sink) if all the three methods identified it as a source (or a sink) and it had at least two closed contours surrounding it. All three methods were implemented in Matlab^®^. The size and strength of a source or a sink was determined from the closest (innermost) contour. The size was defined as the number of pixels inside the contour while the strength was defined as the contour value. The contour borders defined the shape of the source or sink.

#### 2.3.2 Determination of trajectories of pixels and their temporal speeds

In order to determine trajectories of pixels, streamlines were estimated from the velocity vector fields using the built-in “stream3” function of Matlab^®^. A streamline represents the curve to which a subset of velocity vectors in the field are tangent and thus it represents path of the flow of a pixel. The stream3 function calculates a temporal streamline from the velocity vector fields for a pixel of interest defined in space-time. The temporal velocity of the pixel along its trajectory was then calculated from the streamline.

### 2.4 Simulated data and addition of noise

To compare the analytical efficacy of TS, CLG, and HS methods in determining velocity vector fields and source/sink locations, trajectories of pixels and their temporal speeds, simulated data were generated in Matlab^®^ with known shapes of sources and sinks, and motion profiles (velocities and directions of movement). Three image sequences (explained in the results section) were generated; Traveling half-sinusoid plane wave, traveling half-sinusoid circular wave originating at the center (single source), and complex multisource/multisink traveling Gaussian waves. To add noise to the simulated image sequences, the average power of all pixels (over space and time) was calculated. The amplitude of Gaussian white noise added to each pixel was proportional to the average power. The code for generating simulated data and the addition of noise is provided with the toolbox.

### 2.5 Optical imaging - Data acquisition and analysis

#### 2.5.1 Animals and Surgery

Adult (~25 g) male C57BL/6J mice were used for voltage sensitive dye imaging experiments. “Emx-GCaMP6f” animals were used for calcium imaging. The “Emx-GCaMP6” mice produced by crossing homozygous B6.129S2-Emx1 ^tm1(cre)Krj^/J strain (Jax no. 005628) with B6;129S-Gt(ROSA)26Sor ^tm95.1(CAG-GCaMP6f)Hze^/J strain (Jax no. 024105,Ai95) (39). The presence of GCaMP6f expression was determined by genotyping each animal before each surgical procedure. Mice were housed in clear plastic cages in groups of two to five under a 12 h light, 12 h dark cycle. Mice were given ad libitum access to water and standard laboratory mouse diet at all times. The animal protocols were approved by the University of Lethbridge Animal Care Committee and were in accordance with guidelines set forth by the Canadian Council for Animal Care. At approximately 3 months of age, mice were given an acute craniotomy. Mice were anesthetized with isoflurane (1.0–1.5%) for induction and during surgery, and a reduced maintenance concentration of isoflurane (0.5%) was used later during data collection. Mice were placed on a metal plate that could be mounted onto the stage of the upright macroscope, and the skull was fastened to a steel plate. A 7 ×6 mm unilateral craniotomy (bregma 2.5 to –4.5 mm, lateral 0 to 6 mm) was made, and the underlying dura was removed, as described previously (16,40). Throughout surgery and imaging, body temperature was maintained at 37 °C using a heating pad with a feedback thermistor.

#### 2.5.2 Optical imaging

For voltage-sensitive dye based wide-field optical imaging (VSDI), the dye RH-1691 (Optical Imaging, New York, NY) (41) was dissolved in 4-(2-hydroxyethyl)-1-piperazineethanesulfonic acid (HEPES)-buffered saline solution (0.4 mg ml^−1^) and applied to the exposed cortex for 30-60 min, as previously described (42–45). After staining, rest of the dye was washed out and the cortical surface was covered with 1.5% agarose in HEPES-buffered saline and a coverslip on top to reduce movement artifacts. The dye was excited by a red LED (Luxeon K2, 627 nm center) and the excitation and emission optical filters for imaging were 630 ± 15 nm and 688 ± 15 nm (Semrock, New York, NY), respectively. For calcium imaging, mice were anesthetized with isofluorane and their skull was exposed (left hemisphere). Thinning of the skull bone was then performed followed by affixing of a glass coverslip with cyanoacrylate glue and implantation of a metal head plate. The fluorophore GCaMP6f was excited with a blue LED (Luxeon, 470 nm center) filtered by an excitation filter 470 ± 40 nm (Semrock, New York, NY). The emitted light was filtered by a 542 ± 27 nm emission filter (Semrock, New York, NY). 12-bit CCD camera (1M60 Pantera, Dalsa, Waterloo, ON) was used to acquire images at a rate of 150 and 30 frames per second for VSD and calcium imaging. Because animal brain states show spontaneous change, we averaged 10–45 trials of stimulus presentation to reduce these effects. To correct for time-dependent changes in fluorescent signals that accompany all imaging, we also collected a number of nonstimulation trials that were used for normalization of the stimulated data. A 10-s interval between each sensory stimulation was used. The camera was focused 0.5-1 mm below cortical surface to avoid distortion of signal due to movement of superficial blood vessels. The preprocessed data is provided with the toolbox data residing in subfolder “SampleData” in the main toolbox folder.

#### 2.5.3 Sensory stimulation

Sensory forelimb stimulation was done by passing a current (0.2-1 mA for 1 ms) through a thin needle (0.14 mm) inserted into the forepaw. For auditory stimulation, a 12 kHz pure tone at 80 dB was played using a Tucker-Davis Technologies (TDT) RX6 and delivered to animal’s right ear while the animal was sitting in a sound proof booth in the laboratory. The speaker (TDT, ES1 electrostatic loudspeaker) was calibrated to emit uniformly distributed amplitude of all frequencies.

#### 2.5.4 Data Preprocessing

To reduce regional bias in caused by uneven dye staining or brain curvature, the VSD and calcium responses were expressed as a percentage change relative to baseline (∆F/F0× 100%) using Matlab^®^. To reduce the effect of heartbeat in our calcium imaging data, the calcium signal was filtered with a finite-impulse-response low-pass filter (cut-off frequency 5Hz).

#### 2.5.5 Statistical Analysis

All statistical analysis was done in Matlab^®^. Two sample t-test was used for statistical comparison and a p-value of 0.05 was used to determine significance.

## 3. Results

We first present the comparison of the performance of the three optical-flow (OF) methods using simulated data for which the ground true values of wave motion are known. Later, we demonstrate the use of OF methods on real voltage and calcium optical imaging data.

### 3.1 Optical-flow characterization of simulated data

#### 3.1.1 Travelling plane wave (no source and sink): a half-sinusoid moving in one direction

One of the most commonly observed wave pattern in brain activity is a travelling plane wave (3,16,29,46–49) which is encountered when an activity has emerged from a source and is travelling towards a sink. It would also be seen in situations where the imaging field of view is limited. We mimicked such activity by a half-sinusoid travelling plan wave at a constant speed (1 pixel per frame – p/f) in the horizontal direction (angle = 0, see Fig 2A and Supplementary Movie 1).

**Figure 2.**
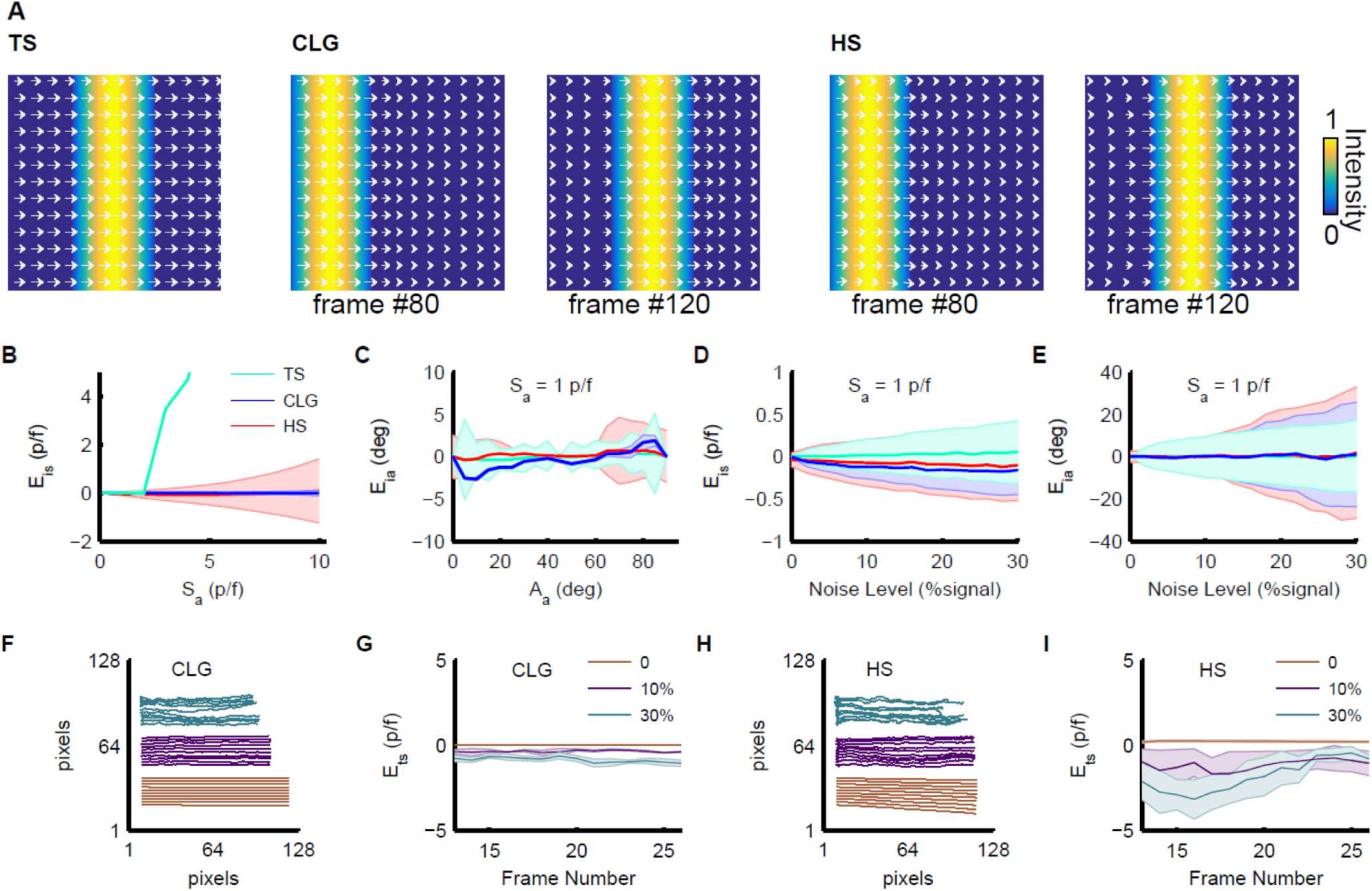
Comparison of the performance of HS, CLG, and TS methods for estimating dynamics of a simulated traveling plane wave. **A**. Representative frames from the image sequence with overlaid velocity vector fields. Note that with the TS method only one vector field is obtained for the whole image sequence while with the HS and CLG methods, a vector field is obtained for each pair of consecutive frames (see text) **B**. Instantaneous speed estimation error versus actual speeds of plane wave for angle = 0°. Performance of the TS method depends on the size of correlation window in space (see text) and shows high error for speeds where displacements of pixels are beyond the size of the correlation window. HS shows an increase in the standard deviation of the error with increasing speeds but CLG is robust. **C**. Instantaneous angle estimation error versus actual angles (direction of motion) of plane wave with speed = 1 p/f. TS method shows large standard deviation of error. HS and CLG methods show similar performance with the CLG method showing smallest standard deviation. **D, E**. Instantaneous speed (D) and angle (E) estimation error versus noise level. The performance for three methods degrades with increasing noise levels. **F-I**. Trajectories of pixels for 0% (orange), 10% (purple) and 30% (green) noise level shown in single frame (F) for CLG, and (H) for HS methods. Temporal speed estimation error versus time (frame number) for the three noise levels (G) for CLG, and (I) for HS methods. Overall the best performance is that of CLG.

##### Determination of velocity vector fields

With the three algorithms (TS, CLG, and HS), optical-flow of the travelling half-sinusoid was determined i.e. velocity vector fields were estimated where velocity vectors obtained for all pixels represent their speeds and directions of motion (see Fig. 2A). Note that with the TS method only one vector field for the whole image sequence representing average speeds and directions is obtained. With the CLG and HS methods however, vector fields are estimated from all pairs of consecutive frames and thus their number is one less than the number of frames in the image sequence. The velocity vectors for the CLG and HS methods thus represent instantaneous speeds and directions. The estimated vector fields with all three methods very well matched with the actual flow (Fig. 2A). Moreover, flow fields calculated with the CLG and HS methods were similar to each other. The time required for the calculation of velocity vector fields was longest for the TS method (~79 secs for estimating 1 velocity vector field from 242 frames each having 128x128 pixels). The time took by the CLG and HS methods were 0.52 secs and 0.59 secs respectively for generating 1 velocity vector field from two consecutive frames. These times were obtained on a 64-bit PC with the following specifications; Processor Intel^®^ Core™ i7-4770K CPU @ 3.50GHz, RAM 32GB, and with toolbox running in Matlab^®^ R2013b.

##### Performance of HS, CLG, and TS methods for determining velocity vector fields

In order to gauge the efficacy of the three methods in determining velocity vector fields (speeds and angles), we compared estimated and actual instantaneous velocity vectors by determining the errors for a range of values of speeds and angles (directions of motion) of the sinusoids:

E_is_ = estimated speed (S_e_) – actual speed (S_a_), and

E_ia_ = estimated angle (A_e_) – actual angle (A_a_)

First, E_is_ was measured for all pixels and the trend of means and standard deviations of E_is_ was calculated for values of actual speeds ranging between 0 and 10 pixels/frame - p/f (Fig. 2B) while keeping the angle = 0 (horizontal direction of motion). The TS method performed well for actual speeds less than 2p/f but its performance steeply declined for greater speeds because of limited size of the time window used for determining temporal correlations (see methods). The TS method is therefore sensitive to the size of window chosen for analysis. A larger window would produce better estimates for higher actual speeds but with an added computational cost (Supplementary Fig. S1B). The performance of CLG and HS methods was better than TS for higher speeds with CLG performing better than HS (small standard deviation for CLG in Fig. 2B). Similarly, E_ia_ was determined for all pixels and the trend of mean and standard deviations of E_ia_ was observed for values of angles ranging between 0 and 90° (Fig. 2C) while keeping the speed constant = 1p/f. E_ia_ remained within ±5° for all the three methods with individual variations in the values of means and standard deviations. Although the mean value of E_ia_ calculated with the TS method is closer to zero as compared to means calculated by using the HS and CLG methods, the standard deviation of E_ia_ calculated with the TS method is much higher. The performance of CLG and HS in estimating the angles is similar with a very small standard deviation of error with the CLG method. Based on these results, the CLG method should be preferred over TS and HS methods for determining velocity vector fields for travelling plane waves.

##### Efficacy of HS, CLG, and TS methods in determining vector fields from noisy data

Since, the experimental data representing brain activity is noisy, we tested the performance of the three optical-flow methods by adding noise to the simulated data (see methods). By adding various levels of noise to each pixel in the image sequence, the errors E_is_ and E_ia_ in speed and angle estimation respectively were calculated in the same way as mentioned above. The addition of noise adversely affected the estimates of speeds and angles with larger errors in estimation for larger noise levels (Fig. 2D and 2E). The estimate of angles however, were more affected with greater noise (Eia ~30° for 30% noise level). From all the three methods, the temporospatial method was more robust in the estimation of angle and velocity from noisy data as compared to the other two methods. This is because determining the temporal correlation between pixels improves the signal to noise ratio. The performance of CLG and HS methods was similar to each other for all noise levels with CLG having smaller standard deviations of errors. Our results thus suggest that filtering imaging data to remove noise is preferable before determining vector fields.

##### Determination of trajectories of pixels and their speeds versus time

To determine underlying anatomical and physiological communication pathways within different brain regions, it is relevant to follow the course of travel of activity. Hence, using vector calculus methods, the trajectories of travel (streamlines) of pixels or regions of interest were determined from velocity vector fields calculated with CLG and HS methods. Note that this analysis cannot be done with the single velocity vector field obtained with the TS method. Estimation of trajectories from velocity vector fields determined with the CLG method were better than that of the HS method (compare red lines in Fig. 2F and 2H). However, when noise (10% and 30%) was added to the simulated data, the performance of estimation declined similarly for both methods. Temporal speeds (S_t_) of pixels along the trajectory were also estimated and the error (E_ts_ = estimated S_t_ – actual S_t_) was calculated. The error was zero for simulated data with no noise but increased for 10% and 30% noise levels (purple and green lines in Fig. 2G and 2I). The error was worse for analysis based on the HS method (Fig. 2I). Note that the displacement (direct distance from start to finish) of a pixel was smaller at higher noise levels. This is mainly because of error in the estimate of angles resulting in zig-zag shaped trajectories of pixels. The zig-zag effect grows larger with increasing noise levels. Our results thus suggest that the trajectories of pixels and their speeds are better estimated from velocity vector fields determined with the CLG method after noise reduction.

#### 3.1.2 Travelling circular wave: a half-sinusoid originating at a source and propagating in all directions

Another frequent pattern of activity in the imaging data is propagation of a signal originating from a source such as those seen after sensory stimulation (16,49–51). In order to mimic this type of activity, a circular ring of half-sinusoid was generated which originated at the center of a frame and spread outwards to the edges with a constant velocity of 1 p/f (Supplementary Movie 1). The optical-flow was estimated with the three methods to obtain velocity vector fields (Fig. 3A). Errors in the estimates of instantaneous speeds and angles (E_is_ and E_ia_) were determined for propagation speeds ranging between 0 and 10 p/f (Fig. 3B-C). Note that in this simulated data all directions of motion i.e. all angles are concurrent and hence only the propagation speed was changed. Similar to the results above, performance of the TS method degraded for estimation of higher speeds (Fig. 3B) because of limited size of window used for finding temporal correlations between pixels (see methods). The TS method also failed to estimate the directions of motion for higher speeds for the same reason (Fig. 3C). The performance of the CLG and HS methods was similar to each other with CLG’s performance better than that of HS for estimating speeds as well as angles (smaller standard deviations of errors for CLG curves).

**Figure 3.**
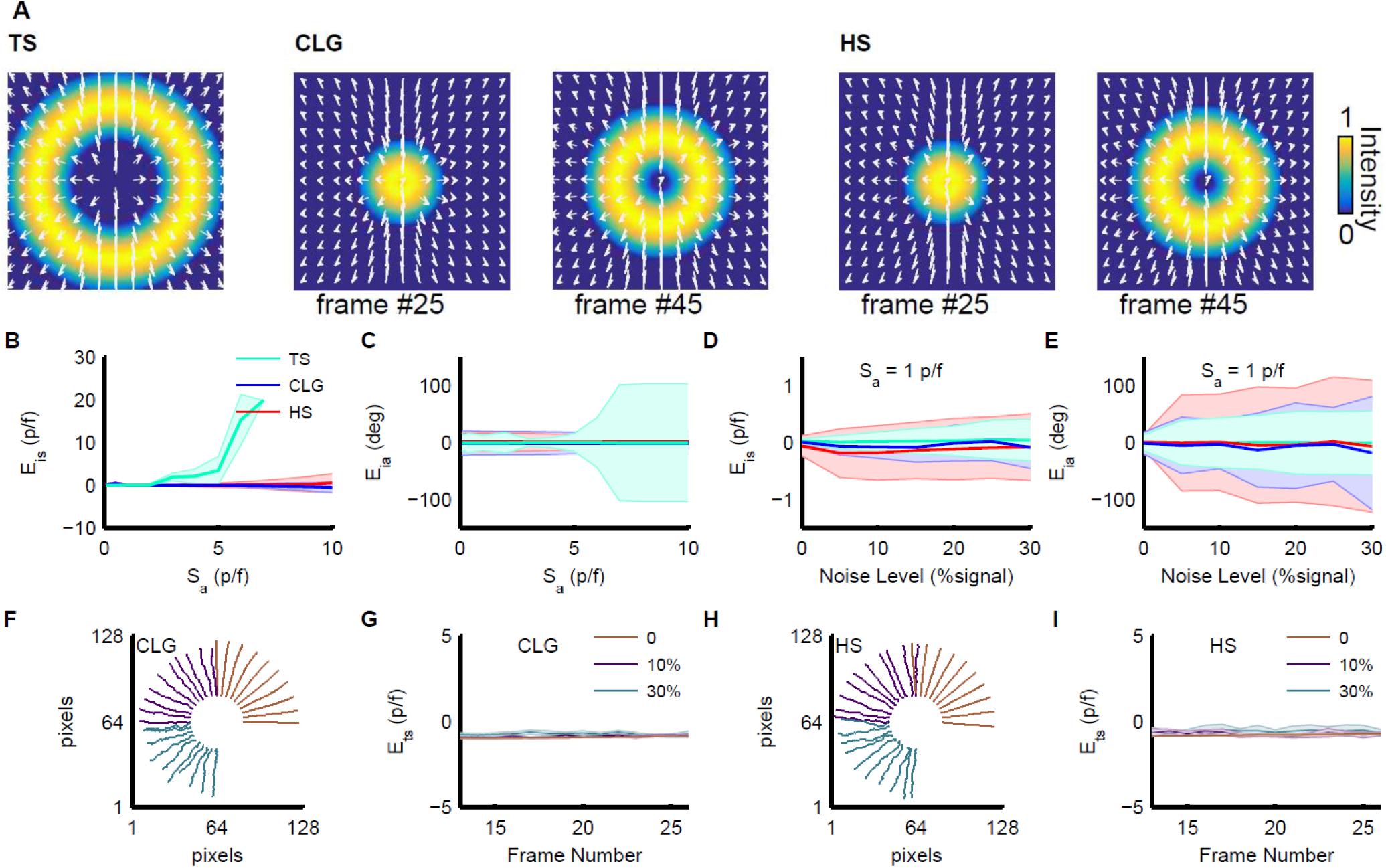
Comparison of performance of HS, CLG, and TS methods for estimating dynamics of a simulated traveling circular wave. **A**. Representative frames from the image sequence with overlaid velocity vector fields. **B, C**. Instantaneous speed estimation error (B) and instantaneous angle estimation error (C) versus actual speeds. **D, E**. Instantaneous speed estimation error (D) and instantaneous angle estimation error (E) versus noise levels. **F-I**. Trajectories of pixels for 0% (orange), 10% (purple) and 30% (green) noise level shown in single frame (F) for CLG, and (H) for HS methods. Temporal speed estimation error versus time (frame number) for the three noise levels (G) for CLG, and (I) for HS methods. Overall the best performance is that of CLG.

The efficacy of the three methods for determining velocity vector fields from noisy data was assessed by adding different noise levels to all pixels and observing the errors in estimates of instantaneous speeds and angles. The propagation speed was set to 1p/f for this analysis. The TS method was most robust to noise as computing temporal correlations averages out some noise (Fig. 3D-E). However, all three methods were better in estimating speeds from noisy data than angles. With the addition of even small noise levels, the performance of the HS method in estimating angles degrades quickly. These results suggest that it is preferable to filter raw data before applying any of the three methods for determining velocity vector fields.

Next, trajectories of selected pixels and their velocities were calculated from velocity vector fields that were estimated with the CLG and HS methods with and without adding noise to all pixels. Without the addition of noise, the estimated trajectories were accurately determined (red lines in Fig. 3F and 3H). However, with the addition of 10% noise level, the estimated trajectory is such that the total displacement of pixels is shorter than the actual (purple lines). This is because the estimation of angles of velocity vector fields is adversely affected with the addition of noise and thus the estimated path is zig-zag. The zig-zag effect becomes more pronounced with the addition of 30% noise level (green lines). The effect of noise on the estimation of temporal speeds was negligible (Fig. 3G and 3I) with the CLG method based analysis showing smaller standard deviations of error. Combined these results also favor the use of CLG method for determining velocity vector fields.

#### 3.1.3 Travelling Gaussian waves: Complex simulated data containing three Gaussian sources, travelling waves, and sinks

When multiple brain regions are active simultaneously (16,42,50,52), multiple travelling waves are observed (see Fig. 1b in (16)). In order to gauge the correctness of our analyses in estimating the spatiotemporal dynamics of activity in such data, we generated similar simulated data by forming an image sequence in which three two-dimensional Gaussian sources (with same amplitude = 1) originate at three distinct space-time locations i.e. the spatial points of origin are distinct as well as the frames in which they start (Supplementary Movie 1). Based on their shapes, these activity-like events were named C1 (circular), E1 (elliptical), and C2 (circular). The temporal order of their appearance was C1, E1, and C2. All events originate and expand to reach their final size and when fully expanded, undergo translation and/or rotation. The trajectories of C1 and E1 form arcs of different circles and the trajectory of C2 is linear. E1 while translating also rotates 30° counter-clockwise about its center. Finally, all of them contract and sink into three distinct space locations. For each event, the full expansions and contractions happen in 15 frames (~0.1 secs). For C1 and E1 during translations while following arcs of circles, the variation in angular speed follows a half-sinusoid while for C2, the linear translation happens at constant speed. By generating this multi-brain activity-like complex simulated data our goal was to determine how accurately our analysis would estimate the underlying dynamics and to compare the performance of CLG and HS methods. The TS method was not used here because it requires large computational time and provides only a single vector field for the whole image sequence.

Velocity vector fields were first estimated (Fig. 4A for the CLG method) and later using methods of vector calculus, the trajectories of pixels of interest and their speeds were calculated (Fig. 4B-I for C1 and E1). The analysis based on the CLG method accurately captured the dynamics of C1 and E1 (with a few exceptions for E1 – see below). The analysis based on the HS method however was not as much efficient. For example, the center of C1 is static while the Gaussian is expanding, follows a half sinusoid speed curve while translating along the arc, and is static again while the Gaussian is contracting (see dashed magenta line for actual speed of center in Fig. 4C). The estimated speed of the center of C1 with the CLG method based analysis closely matched the actual speed (see green lines in Fig. 4B and 4C for trajectory and speed respectively). However, the HS method based analysis failed to capture the dynamics accurately during the expansion and contraction phases (green lines in Fig. 4D-E). For non-center points in C1, after their temporal appearance, the actual speed is ~2p/f while C1 is expanding, follows a half sinusoid speed curve (linear speed = radius of point x angular speed), and is ~2p/f while C1 is contracting. The estimated speeds of these non-center points also closely matched the actual speeds (see blue, black, and red lines in Fig. 4B-C for CLG method based analysis). The HS method based analysis fails to accurately capture the dynamics of non-center points (Fig. 4D-E). Note, that there is a momentary pause of C1 Gaussian for two frames between expansion and translation, and for two frames between translation and contraction. This is also nicely captured in the estimates of speeds of non-center pixels of C1 especially with the CLG method based analysis.

**Figure 4.**
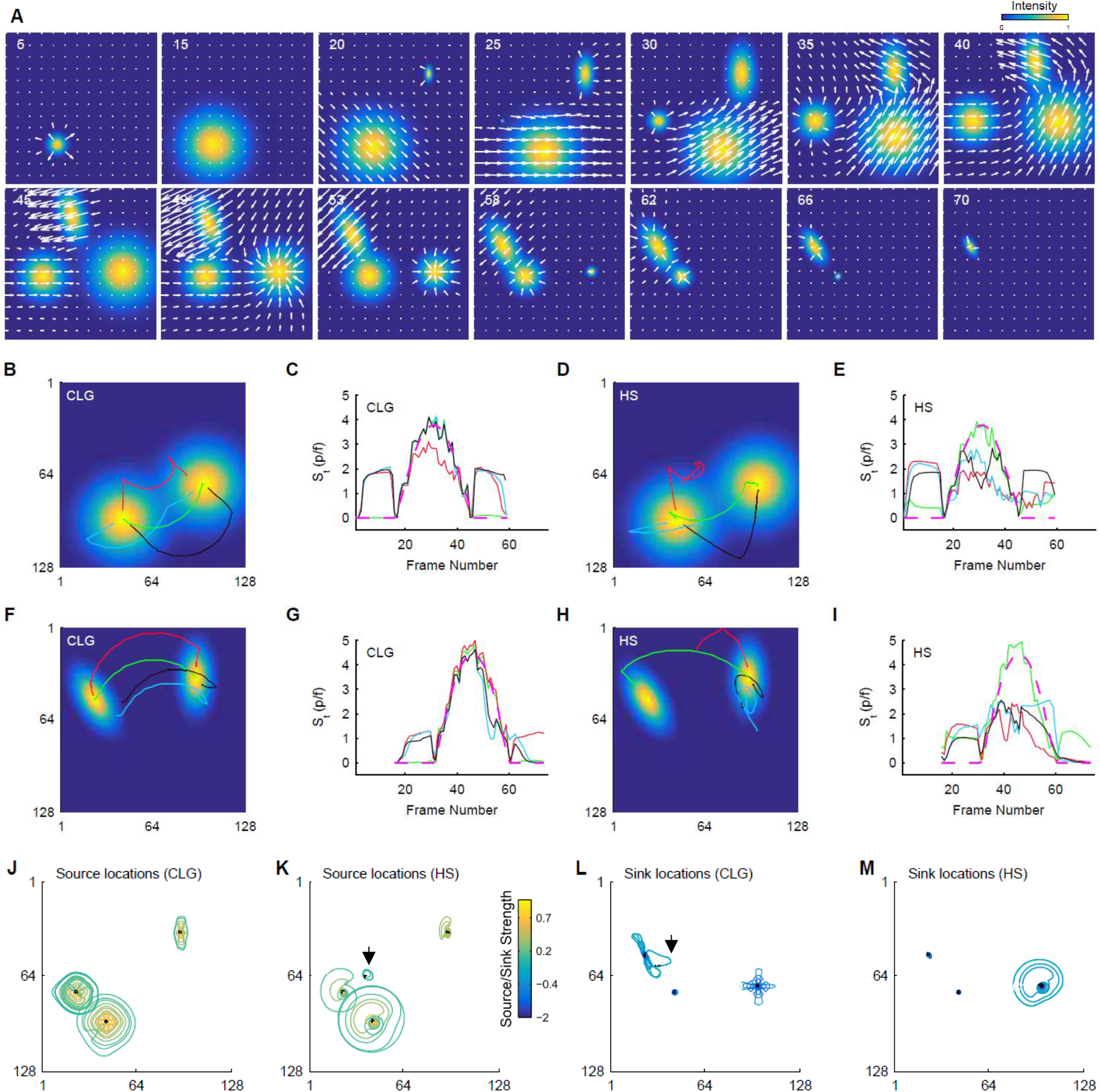
Comparison of the performance of HS and CLG methods for estimating dynamics of an image sequence simulating three sources, traveling events, and sinks (see text). **A**. Montage of frames from the image sequence overlaid with velocity vector fields determined with the CLG method. **B**. Trajectories of selected pixels of C1 event (see text), center pixel (green), non-center pixels (blue, black, and red), calculated with the CLG method based analysis. **C**. Estimated temporal speeds of pixels in (B) - same colors, versus time (frame numbers). Magenta line shows actual speed of the center pixel. **D-E**. Same as B-C but with the HS method based analysis. **F-G, and H-I** Same as B-C and D-E respectively for E1 event. **J-K**. Estimation of source locations (black dots) and shapes/outlines (contours) with the CLG and HS methods. **L-M**. Estimation of sink locations with the CLG and HS methods. Contour colors show source/sink strengths. The CLG method wins in estimating pixel trajectories and temporal speeds and works favorably for source/sink analyses.

For E1 also, the estimated trajectory and speed of the center pixel (green line in Fig. 4F-G) closely matched the actual trajectory and speed (dotted magenta line) respectively with the CLG method based analysis. However, the estimated trajectories and speeds were slightly erroneous for non-center pixels (see black, blue, and red lines in Fig. 4F-G). This error could in part be due to discrete spatial sampling and rotation of E1 event during translation. Here also, the analysis captures the momentary pause before translation along the arc. With the HS method based analysis, the estimation of event dynamics (trajectories and speeds) was not as accurate (Fig. 4H-I). These results thus demonstrate the superiority of the CLG method based analysis over that of HS in tracking pixel trajectories and determining temporal speeds.

Since simulated data had sources and sinks associated with three simulated activity patterns, we applied vector calculus methods on each velocity vector field determined by the CLG and HS methods to identify the location, outlines, and strengths of sources and sinks in space-time (see methods). Both CLG and HS method based analyses performed well in identifying the location of sources and sinks (see black dots in Fig. 4J-M). Although there were only three events, but since the outflow of activity from a region is identified as a source, multiple sources were detected in space-time where C1, E1, and C2 were detected as expanding. Similarly, multiple sinks were detected. Most of the identified source and sink locations except a few, were very close (within a few pixels) to the actual locations. A weak source (relatively smaller contour value) identified with the HS method based analysis and a weak sink identified with the CLG method based analysis were a little farther from the actual locations (arrows in Fig. 4K and 4L). These fake sources and sinks are perceived in regions where events are spatially close and move away or towards each other respectively. Hence, caution is advised while interpreting whether a detected location is a real source or a sink when multiple events are present and approach or move away from each other in experimental data.

The characteristics of sources and sinks i.e. their outlines (shapes), sizes, and strengths were also estimated by finding the contours from the divergence of vector fields (see methods). The size of a source (or a sink) was defined as the area of the contour detected around it while the contour’s value (divergence) was its strength. Both CLG and HS methods based analyses identified the outlines (contours) of sources and sinks with the former performing better qualitatively (see contours in Fig. 4J-M). With both types of analyses, the stronger sources (yellower contours in Fig. 4J-K) and sinks (bluer contours in Fig. 4L-M) were correctly detected close to the center location. Additionally, larger sizes were detected gradually with increasing time (frame numbers) as C1, E1, and C2 expanded. We also observed the product of size and strength (strength weighed size or vice versa) as a useful parameter to determine the significance of a source or sink i.e. larger product would indicate a significant source or sink activity (data not shown). The quantification of parameters for sources and sinks discussed here will be useful for pin pointing the origin and termination of activity spread in real image sequences as discussed below in section 3.2.1.

##### Limitations of the CLG method based analysis in estimating trajectories and speeds of pixels; a word of caution

The CLG method based analysis failed to accurately estimate the dynamics of pixels which were shared by multiple dynamic events in space-time. For example, the C2 activity starts and ends in frames 24 and 68 of the image sequence respectively (see Supplementry Movie 1). The estimated trajectories and speeds of all C2 pixels between these frames were erroneous (representative pixels in Fig. 5A-B). The actual speeds of all C2 pixels except the center were ~1.43p/f during expansion and contraction with a momentary pause and linear translation at 1.3p/f in between. The center pixel only moves during translation. The estimated trajectories and speeds however, were nowhere close to the actual ones because the existence of C2 wave in frame 24 is concurrent and within the boundaries of C1 event (see Supplementary Movie 1). The CLG method based analysis thus captures the motion dynamics of the pixels of C1 and continue to follow its wake in subsequent frames. For the center pixel of C2, the estimated trajectory is that of pixel of C1. When the center C2 pixel appears in time (in frame 24), it causes a discontinuity in the flow field which is ignored by the CLG method due to the spatial smoothness constraint (Eq. 2 – see methods) and perhaps due to the motion constancy constraint as well (Eq. 4). When the C2 was estimated starting frame 29 after coming out of the wake of C1, the trajectories and speeds of both center and non-center pixels were accurate compared to previous values (Fig. 5C-D) with the exception of some pixels which towards the end entered into the wake of E1 activity and thus the estimates of their trajectories and speeds were erroneous (see blue line in Fig. 5C-D). These results suggest that if events are overlapped in space-time, the estimation of trajectories and speeds can be erroneous and caution has to be taken in identifying the pixels of interest in both space and time.

**Figure 5.**
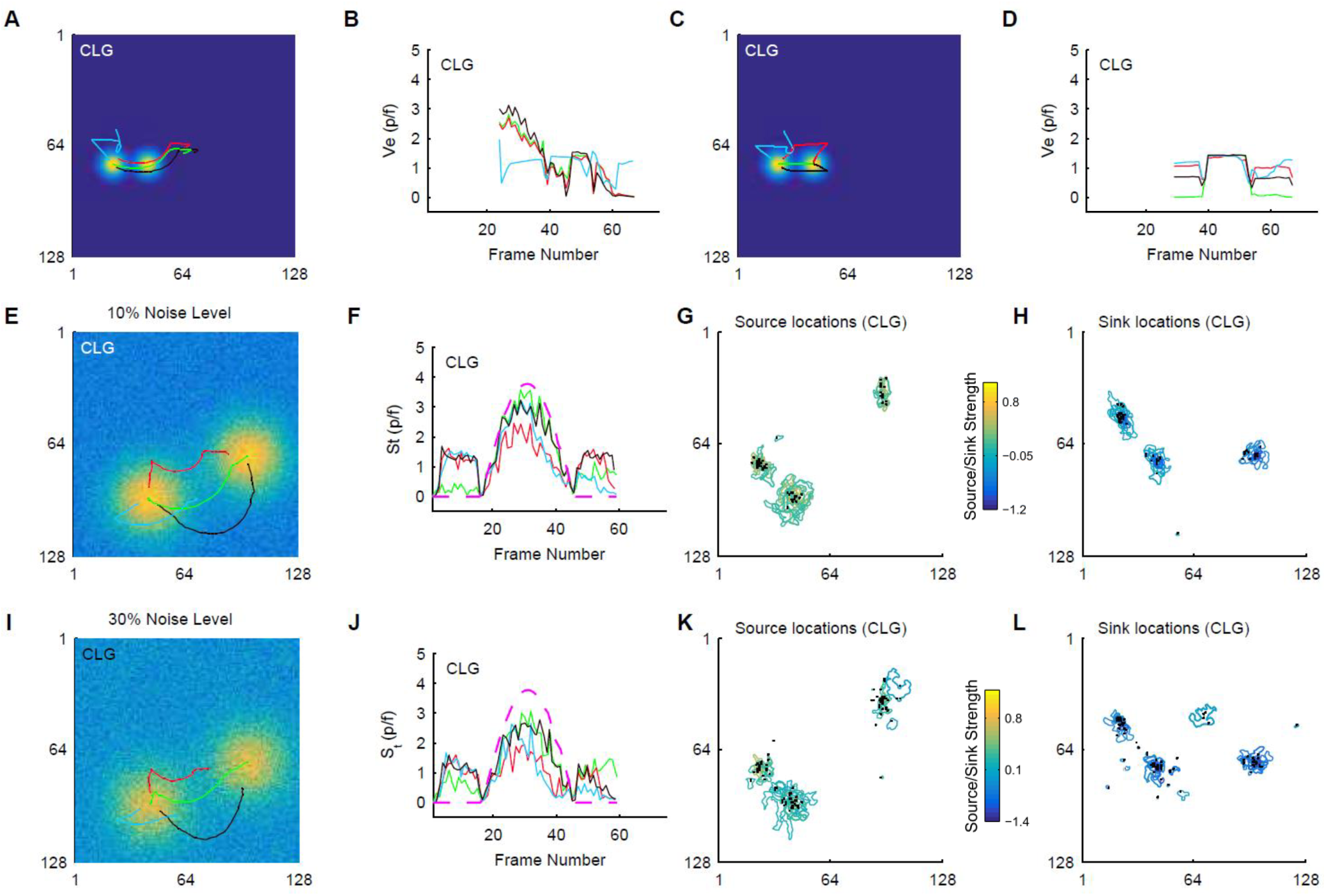
Limitations of the CLG method based analysis – errors due to the addition of noise and/or overlap of simulated events in space-time. **A**. Erroneous estimation of trajectories of selected pixels of C2 starting frame 24 because at this time, C2 is in the wake of C1 (see text). Center pixel (green), non-center pixels (blue, black, and red). **B**. Erroneous estimated temporal speeds of pixels in (A) - same colors, versus time (frame numbers). **C-D**. Same as B-C but correct estimation of trajectories and temporal speeds with analysis starting frame 29. **E-H,** Same as Figure 4 B-C, J, & L but with 10% noise level added to the image sequence. **I-L**. Same as E-H with 30% noise level added to the image sequence. The addition of noise reduces the performance of the CLG method. Caution is advised for selecting pixels and regions of interest in space-time for determining trajectories and temporal speeds.

We further tested the errors in optical-flow estimation (with both CLG and HS methods) due to overlapping events by changing the amplitude of individual Gaussian events (C1 = 2.5, E1 = 10, and C2 = 5; see supplementary movie 1 and Fig. S2). With these values of amplitudes, there is a larger spatiotemporal overlap between the values of three motion patterns. When E1, C1, and C2 are concurrent in a frame, farther away from E1’s center, its pixel values would be close to the amplitudes of C1 and C2. New position of an overlapping pixel in the next frame hence will not be estimated accurately. With executing optical-flow analyses on this dataset, the performance of both CLG and HS methods degraded. However, the CLG method based analyses still performed better than the HS method in determining pixel trajectories and temporal speeds (compare Fig. S2 B,C and F,G with D,E and H,I respectively). The locations of sources determined from velocity vector fields estimated with the CLG method were qualitatively better than those estimated with the HS method while the opposite was true for determining sink locations (Fig. S2 J-M).

##### Limitations of the CLG method based analysis - sensitivity to the addition of noise

The efficacy of the CLG method based analysis in determining velocity vector fields, trajectories of pixels and temporal speeds from noisy data was assessed by adding 10% and 30% noise levels to all pixels. The addition of noise adversely affected the estimates of trajectories and speeds of pixels and identification of the location of sources and sinks with larger effects observed for larger noise levels (Fig. 5E-L shown only for C1; compare with Fig. 4B-C, 4J, and 4L). For example, the trajectory of the center pixel (green line in Fig. 5E and 5I) was not estimated to be purely arc of a circle but has a translation component as well. For non-center pixels, the estimated trajectories are incomplete (red, blue, and black lines) with dislocated final pixel points. Similarly the estimated speed is erroneous for center and non-center pixels (Fig. 5F & 5J) but the overall trend of motion was successfully captured i.e. expansion, translation, and contraction. These errors in the estimates of pixel trajectories and temporal speeds are due to errors in the estimates of velocity vector fields by the CLG method i.e. the errors in the estimates of instantaneous speeds and angles (see Fig. 2B-E and 3B-E). The estimates of characteristics of sources and sinks were also adversely affected e.g. multiple sources and sinks are estimated near the actual source/sink points (Fig. 5G-H & 5K-L). However, non-existing sources and sinks that were estimated were spatiotemporally located very close to the actual space-time locations. Estimates of outlines of sources and sinks were also adversely affected with the addition of noise however, the strengths were still the highest near the origin point of sources and sinks (yellower and bluer contours respectively near the center). These results thus suggest that noise adversely affects the analysis and filtering of raw data is advisable before determining velocity vector fields.

### 3.2 Optical-flow characterization of real optical imaging data

#### 3.2.1 Wide-field optical imaging of mouse brain with forelimb sensory stimulation

Wide-field imaging of mouse brain was done using voltage-sensitive dye while mouse forepaw was electrically stimulated. An increase in the intensity of the fluorescence signal was observed 10-15 ms after forelimb stimulation in the primary forelimb (FLS1) area of the sensory cortex (Fig. 6A and supplementary movie 2). The activity expanded initially after appearing and translated towards the medio-caudal direction. Sequentially, another activity appeared in the secondary forelimb sensory area (FLS2) smaller in size and with small translation towards the medio-caudal direction. Two regions of interest (ROIs) were defined on the primary and secondary forelimb cortical areas (Fig. 6B) after visual inspection of the image sequence. The average intensity of pixels in the ROIs sharply increased around frame 34 and then decreased in two steps (Fig. 6C). First, a sharp decrease to a value greater than half of the peak value was observed and then a gradual decrease to the baseline. This is in accordance with previously reported trends of FLS1 and FLS2 evoked responses to forelimb stimulation in isofluorane anesthetized mice (16,43).

**Figure 6.**
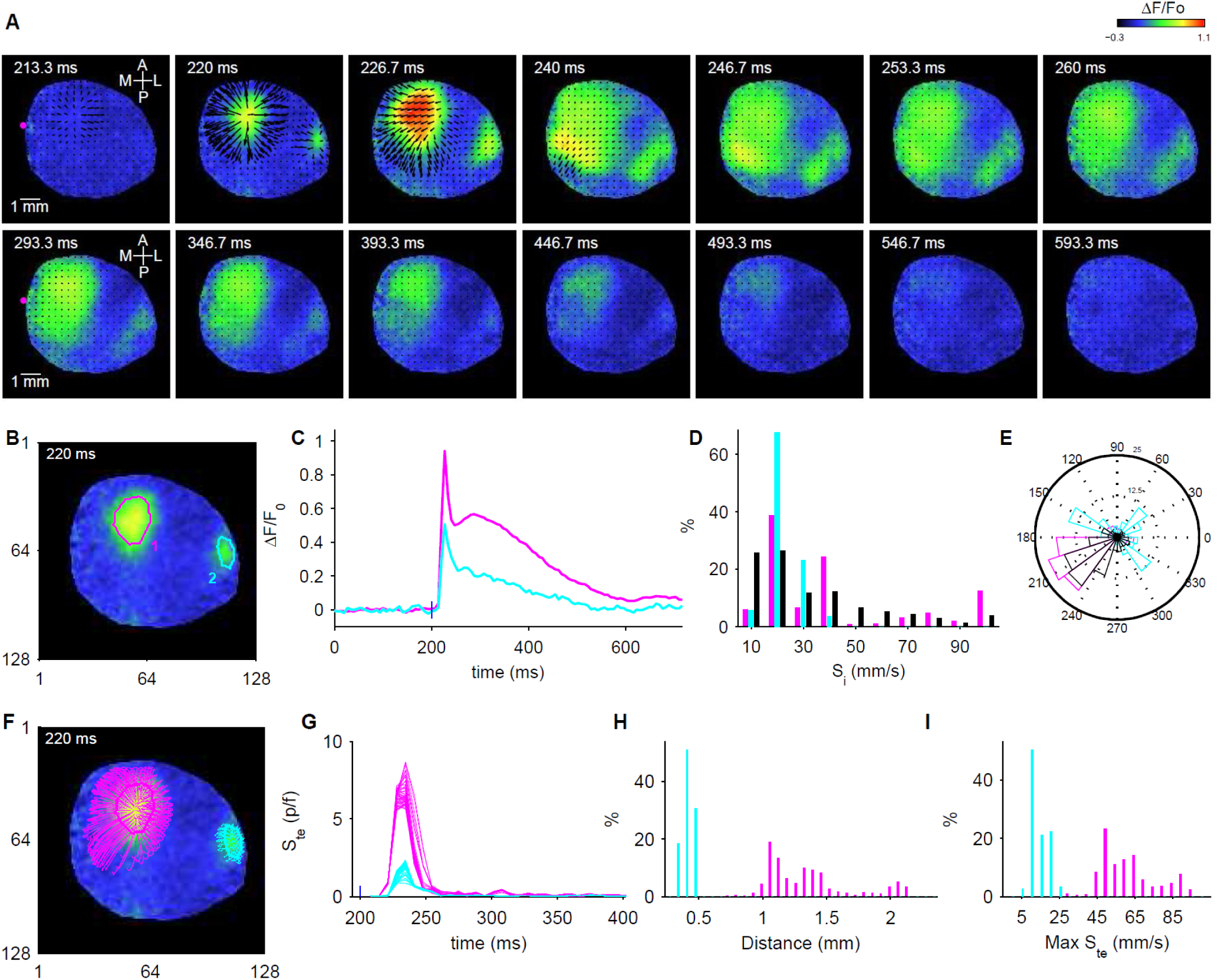
Optical flow characterization (with the CLG method) of forelimb stimulation-evoked VSD activation in anesthetized mouse. **A**. Montage of selected frames from the image sequence with overlaid velocity vector fields. Magenta dot in the first (top left) frame indicates bregma location. **B**. Regions of interest (ROIs) on the primary (magenta) and secondary (cyan) forelimb areas. **C**. ΔF/Fo vs time for ROIs in (B). **D**. Distribution of instantaneous speeds of all pixels in space-time in the ROIs in B (same colors) and for all pixels within the mask (black bars). **E**. Similar to (D) but distribution of instantaneous angles of all pixels within ROIs or the mask. **F**. Trajectories of pixels (shown only for some pixels) in ROIs from (B) – same colors as in (B). **G**. Estimated temporal speeds of all pixels in the ROIs. **H**. Distribution of lengths of trajectories. **I**. Distribution of maximum of temporal speeds. Blue lines in C and G indicate the stimulus onset.

##### Determination of velocity vector fields

The CLG method was used to estimate velocity vector fields of the image sequence, which captured the flow of activity spread (montage Fig. 6A). To analyze the global dynamics of pixel intensity changes in the whole imaged region, instantaneous speeds and angles were determined from velocity vectors for all pixels (in space and time). The distribution of these instantaneous speeds (black bars in Fig. 6D) showed a peak around 20 mm/sec (10 mm/sec = 1 p/f) with a mean ±std value of 33.3 ±24.7 mm/sec and some pixels (~5%) moving as fast as 100 mm/sec. The distribution of angles showed a peak in the medio-caudal direction (black line in Fig. 6E) which is in accordance with the visually observed features of the largest activity in the FLS1 area. Further, analysis was done for FLS1 and FLS2 ROIs to observe local dynamics and compare these functionally connected regions. The distributions of speeds indicated that the activity propagation was faster in the primary as compared to the secondary forelimb area (compare cyan and magenta in Fig. 6D). Mean±std values of speeds for FLS1 and FLS2 ROIs were 42 ± 32.3 mm/sec and 23.2 ± 6.2 mm/sec respectively and the two distributions were significantly different from each other (*** p<0.001) The distribution of angles for the FLS1 ROI showed the largest peak in the medio-caudal direction (Fig. 6E) whereas the FLS2 ROI did not prefer this direction.

Next, to determine the flow of activity of individual pixels, trajectories of pixels, their speeds over time, and lengths of trajectories were calculated for all pixels in both FLS1 and FLS2 ROIs (Fig. 6G-I). Pixels in the FLS1 ROI covered longer distances at higher speeds as compared to pixels in the FLS2 ROI. However, the distribution of distances traveled by pixels in the FLS1 ROI was bimodal indicating two peaks at 1.3 mm and 1.67 mm. As can be seen from the trajectories, the longer distances covered were in the medio-caudal direction (Fig. 6F) whereas the smaller travel was in the opposite and orthogonal directions. The distribution of the maximum value of speeds (Fig. 6I) indicated that most of the pixels in the FLS1 and FLS2 ROIs traveled at speeds of 50 mm/s and 20 mm/s respectively, which agrees with visually observed values.

##### Identification of sources and sinks

Using vector calculus methods, the locations of sources and sinks both in space and time were determined from the velocity vector fields (Fig. 7A-B). Their strengths and sizes were also calculated from contours. Significant sources i.e. ones generating observable activity, were identified by calculating the product of sizes and strengths. Three noteworthy sources (1, 3, and 4 in Fig. 7C) were further analyzed which collocated in space and time with the FLS1 and FLS2 ROIs selected in the analysis shown above (see Fig. 7E-F). For these sources, pixel trajectories for pixels within the contours were determined and their speeds as well as distances covered were determined. The distributions of distances (Fig. 7H) and max speeds (not shown) were very similar to ones obtained with the manual analysis.

**Figure 7.**
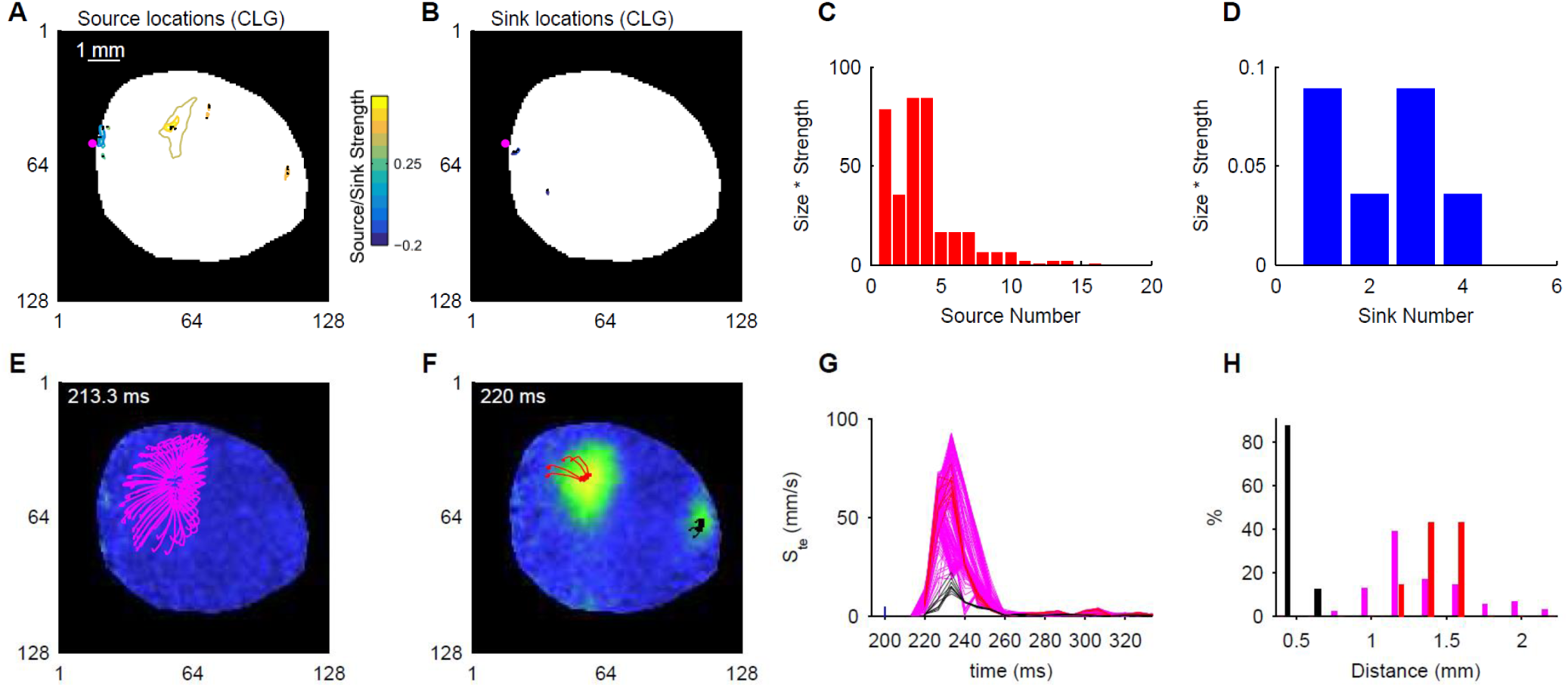
Semi-automatic optical flow analysis (with CLG method). **A-B**. Identified source (A) and sink (B) locations (black dots) and associated contours of VSD imaging data presented in Figure 6. Contour colors indicate strength of sources and sinks. **C**. Size times Strength versus source number for identifying significant sources. **D**. Same as (C) for sinks. **E-F**. Pixel trajectories shown for three selected sources (1 - magenta, 2 - red, and 3 - black) with high size x strength values. Note that source 1 was identified in 213.3 ms whereas sources 2 and 3 were identified in 220 ms. **G-H**. Distribution of temporal speeds (G) and length of trajectories (H) of pixels of sources 1, 2, and 3 (see E-F). The blue line in G shows the stimulus onset. The results here are comparable with those obtained using the manual analysis shown in Fig. 6.

##### Validation of Analysis

The CLG method based analysis allows the study of spatiotemporal dynamics of brain activity sampled with voltage sensitive dye optical imaging. However, since the true velocity vector fields are unknown, we validated the values of speeds and angles of some randomly chosen pixels by determining them manually. Manual validation of results for the FLS1 activity was satisfactory as pixel trajectories and temporal speeds were faithfully estimated. However, the estimated pixel trajectories seemed to be smaller than actual for the FLS2 activity perhaps because of overlapping smaller activities within the same region and smaller sampling rate i.e. discretization in time.

#### 3.2.2 Quantitative comparison of voltage and calcium imaging data

To further validate the efficacy of our analysis, the spatiotemporal dynamics estimated from calcium and voltage imaging were compared (see methods for VSD and Ca imaging with GCaMP6f). With the assumption that the underlying biological activity in the brain is similar for mice stimulated with similar tone stimulus (i.e. frequency and amplitude), we hypothesized that the imaged data using different fluorophores would capture similar spatial dynamics from the auditory cortical area with differences originating from the temporal response dynamics of fluorophores. Since the temporal dynamics of the VSD signal is much faster than that of the GCaMP6f fluorescent protein, we expected estimation of faster velocities in VSD compared to GCaMP6f signal from the auditory cortex (9,36).

After collecting image sequences with the two methods, preprocessing was done similarly to obtain the changes in fluorescence from the baseline (ΔF/F_o_). For both datasets, cortical responses were observed in the primary auditory cortex (AC) and other cortical areas as well. In both image sequences, the latency of response in the AC area was similar (10-20 msec for VSDI and 20-30 msec for Ca imaging). Both activity expanded from primary auditory cortex and after some translation started to sink in the same area. Velocity vector fields for both image sequences were obtained as the output of optical flow analysis with the CLG method. To compare the instantaneous speeds and angles for pixels in the auditory cortical area of the two image sequences, we manually chose regions encompassing the auditory cortex (Fig. 8A-B). As expected, higher instantaneous speeds (Fig. 8C) were sampled with the VSDI (red) as compared to calcium imaging (blue). The distributions of angles (Fig. 8D) qualitatively were different but the largest peaks are in the mediocaudal direction (fourth quadrant) suggesting that the largest flow of activity was comparable in the two datasets.

**Figure 8.**
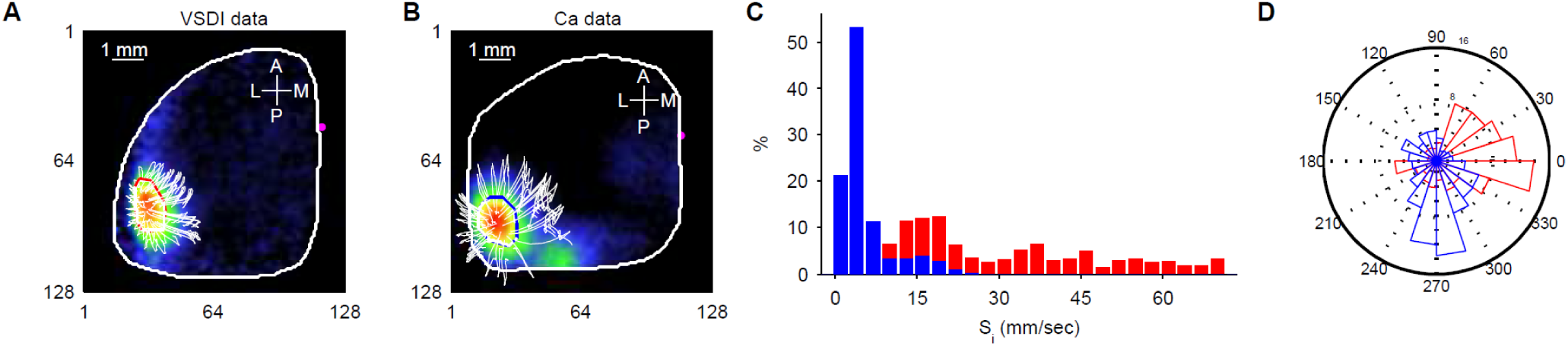
Quantitative comparison of voltage and calcium imaging data. **A**-**B**. Representative images showing extent of activity over the auditory cortical area sampled with VSD (A) and calcium (B) imaging. White lines show travel trajectories for some pixels. Magenta dots indicate bregma location. **C.** Distribution of instantaneous speeds from all pixels within the regions of interest shown in A (red) and B (blue). **D**. Same as C but distribution of instantaneous angles.

### 3.3 **O**ptical-**F**low **A**nalysis Toolbox in **M**atlab^®^ for investigating the spatiotemporal dynamics of **M**esoscale brain activity (**OFAMM**)

We present version 1.0 of the OFAMM toolbox for the analysis of brain activity to be freely used by the research community under the GNU General Public License, version 3 (GPL-3.0) (53). The toolbox and sample data including simulated and real optical imaging data can be downloaded from [<u>http://lethbridgebraindynamics.com/ofamm/</u>]. The toolbox contains a graphical user interface (GUI) shown in Figure 9. This GUI follows the optical brain activity data analysis framework described in Figure 1. The detailed procedure to use the toolbox is provided in the supplementary material. Briefly, one can load image sequence and mask (if applicable) into the GUI. On the image sequence, all or any of the three optical flow methods, HS, CLG, and TS, can be executed to estimate velocity vector fields which can be viewed in the GUI. The distributions of instantaneous speeds of all pixels in space-time within a region of interest can be plotted. Source/sink locations and trajectories of pixels of interest can then be obtained from the velocity vector fields by applying vector calculus analyses. Graphs of source/sink locations and those of pixel trajectories and speeds can be plotted. All the results are stored in mat files and the user can plot additional graphs at will (see supplementary material).

**Figure 9.**
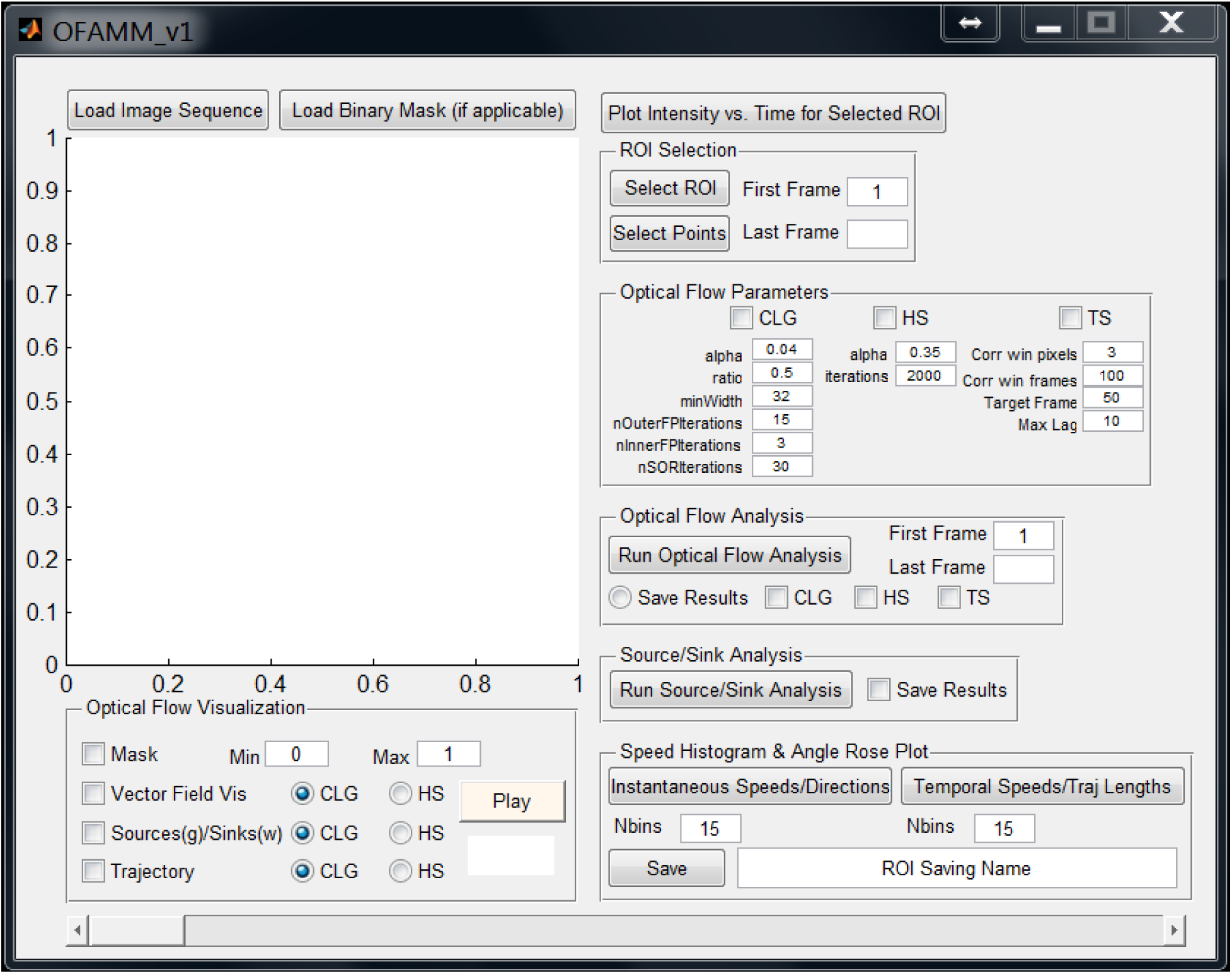
Graphical user interface of “**O**ptical **F**low **A**nalysis Toolbox in **M**atlab for **M**esoscale brain activity” (OFAMM). The sequence of operation of the toolbox for analyzing optical imaging data parallels the flow chart of data analysis shown in Figure 1 i.e. first finding velocity vector fields and then identifying sources, sinks, trajectories of regions or pixels of interest, and their temporal speeds.

## 4. Discussion

Brain activity sampled with optical imaging consists of travelling waves which originate at sources and terminate at sinks. These waves are perceived in sampled image sequences as neurons in different brain regions become sequentially active (3,16,46,54). It is physiologically very relevant to quantify characteristics of such waves, sources, and sinks that might be used to specify the functionality of a brain area and its connectivity with other brain areas (16,17,27,29,55,56) during different brain states and various health status (43).

It is known that many brain disorders result in part from circuit malfunction. The impairment developed in neural circuits and network activity cause instability in neural communication (57–61) which might increase or decrease the cortical dynamics of signal flow or due to circuit remodeling displace the location of sources or sinks. Thus, using these characteristics, normal and diseased brain circuits and their connectivity could be compared (62–64).

Different methods used for the analysis of brain activity in optical imaging datasets have been reported previously but a consolidated toolbox for analyses is missing. Here we present a Matlab^®^ based software toolbox for the optical flow analyses of brain activity datasets. We have incorporated three optical flow methods into the toolbox namely HS, CLG, and TS. Using all three optical flow methods, we have estimated the characteristics of traveling waves, sources, and sinks in brain activity dynamics.

To compare their performance, we generated simulated data with known speeds and directions of movement (data which other investigators can use). The performance of HS, CLG, and TS for determining velocity vector fields was assessed by comparing the estimation errors in speeds and angles without and with the addition of noise (2^nd^ row in Fig. 2 and 3). The results suggest that for noisy data the TS method is quite robust. However, since the TS method relies on determining temporal correlations in a group of frames, only one vector field is obtained where each vector represents average speed and direction. From this single vector field, the location of sources and sinks are determined only in space and not in time. The computational time for executing TS method is also large (Supplementary Fig. S1). In contrast, HS and CLG methods are computationally fast and provide vector fields for pairs of consecutive frames and thus instantaneous speeds and directions. From multiple vector fields location of sources and sinks can be determined in space as well as time. The trajectories of pixels and their temporal speeds can also be determined. For the complex simulated data containing Gaussian propagating waves, we thus compare the performance of HS and CLG methods for determining activity trajectories, their temporal speeds, sources, and sinks. We demonstrate that the CLG method based analyses provide favorable estimates as compared to the analyses based on the HS method (Fig. 4). This is in agreement with previously reported studies where the CLG method performed better in estimating speed of large moving objects in videos and displacement of fluorescently labeled proteins (65,66). For this reason, we perform the analysis of real imaging data only with the CLG method. The user of the toolbox is however encouraged to experiment with all three optical flow methods for their simulated as well as real data.

The parameters for applying optical-flow methods e.g. window size for the TS method, and α and number of iterations for HS and CLG methods can also be set via the graphical user interface before executing the algorithms on an image sequence. Different values of these parameters could affect the estimated velocity vector fields, source and sink locations, and calculated pixel trajectories as well as temporal speeds. For our simulated and real optical imaging data, we also used alternate values of parameters (see methods) to compare their performance (see Supplementary Figures S3-S7). With the use of alternate parameters, the best parameter that improved the performance of HS and CLG algorithms were found (e.g. compare Fig. S5 with Fig. 4). The same conclusions were still valid that (1) the CLG method performs better than HS and (2) filtering data before using optical-flow algorithms is desirable. Since the application of optical-flow methods is sensitive to the choice of algorithm and associated parameters whose values can be set subjectively, we recommend that manual validation of results must be performed.

The CLG method, although is the most favorable, has few limitations. When two or more events overlap in space-time, the CLG method provides erroneous results for the estimation of trajectories of pixels as well as their speeds (Fig. 5 and S2). The errors in estimation were worse with the HS method (data not shown). Caution is thus advised in selecting the pixels or regions of interest in space-time to correctly trace the pathways of activity patterns. Additionally, when noise was added to the signal, the errors in estimation with the CLG method based analysis increased. Thus it is very important to filter the real imaging data before performing the CLG method based analysis.

We demonstrate how the CLG optical-flow method can be applied to real optical imaging data and the distributions of instantaneous speeds and angles, temporal speeds and lengths of trajectories are useful observations for studying population based cortical activity (Fig. 6). Additionally, with the semi-automatic analysis significant sources can be identified and one can observe their spatial distributions (Fig. 7). We validate our optical flow analysis by using two sets of imaging data collected from two different animals with the same stimulation protocol but different fluorophores (VSD and GCaMP6f) for imaging (Fig. 8). As expected, higher instantaneous speeds were captured with VSD as compared to GCaMP6f since the response characteristics of VSD are much faster than GCaMP6f. Finally we present the optical flow analysis toolbox for characterizing the spatiotemporal dynamics in brain activity measured using optical imaging methods (Fig. 9).

Neuroscience relies on observing brain activity at various levels to make physiological models of brain function. At the mesoscale level corresponding to spatial dimensions of the order of hundreds of micrometers to a few millimeters, brain activity is sampled using methods like extrinsic or intrinsic optical imaging. We present a toolbox “**OFAMM**” for analyzing optical imaging data to quantify wave motion and underlying spatiotemporal dynamics.

## Acknowledgements

This work was supported by a Natural Sciences and Engineering Research Council of Canada (NSERC) Discovery Grant #40352, Campus Alberta for Innovation Program Chair, Alberta Alzheimer Research Program to MHM and NSERC CREATE in BIF doctoral fellowship to NA. We thank Jianjun Sun for assistance with surgeries, Behroo Mirzaagha and Di Shao for husbandry, and Jeff LeDue and Tim Murphy for their helpful comments and discussion during the early phase of this work.

